# Curvature generation and engineering principles from *Shewanella oneidensis* multi-flagellin flagellum

**DOI:** 10.1101/2025.02.07.637127

**Authors:** Qing Lou, Hongcheng Fan, Yang Liu, Jeff F. Miller, Yu Huang, Z. Hong Zhou

## Abstract

Motility driven by nanoscale flagella is vital to microbial survival and spread in fluid and structured environments. Absence of native flagellum structures, however, has limited our understanding of the mechanisms of microbial motility, hindering efforts to engineer microbe-based microbots for applications. Here, by cryogenic electron tomography (cryoET) and microscopy (cryoEM), we determined the structural basis of motility driven by the single flagellum anchored to one pole of *Shewanella oneidensis* MR-1 (*S. oneidensis*), an electrogenic bacterium commonly used in biotechnology. The structures of the curved flagellum, representing the conformation during motion, are captured, allowing delineation of molecular interactions among the subunits of its three components—filament, hook, and hook-filament junction. The structures of the filament, i.e., the propeller, reveal a varying composition of the flagellin isoforms FlaA and FlaB throughout the filament. Distinct inter-subunit interactions are identified at residues 129 and 134, which are the major determinants of functional differences in motility for the two isoforms. The hook—the universal joint—has a significantly larger curvature than that of the filament, despite both containing 11 curvature-defining conformers of their subunits. Transition between the propeller and universal joint is mediated by hook-filament junction, composed of 11 subunits of FlgK and FlgL, reconciling incompatibility between the filament and hook. Correlating these compositional and structural transitions with varying levels of curvature in flagellar segments reveals molecular mechanism enabling propulsive motility. Mechanistic understandings from *S. oneidensis* suggest engineering principles for nanoscale biomimetic systems.

**Graphic abstract:** 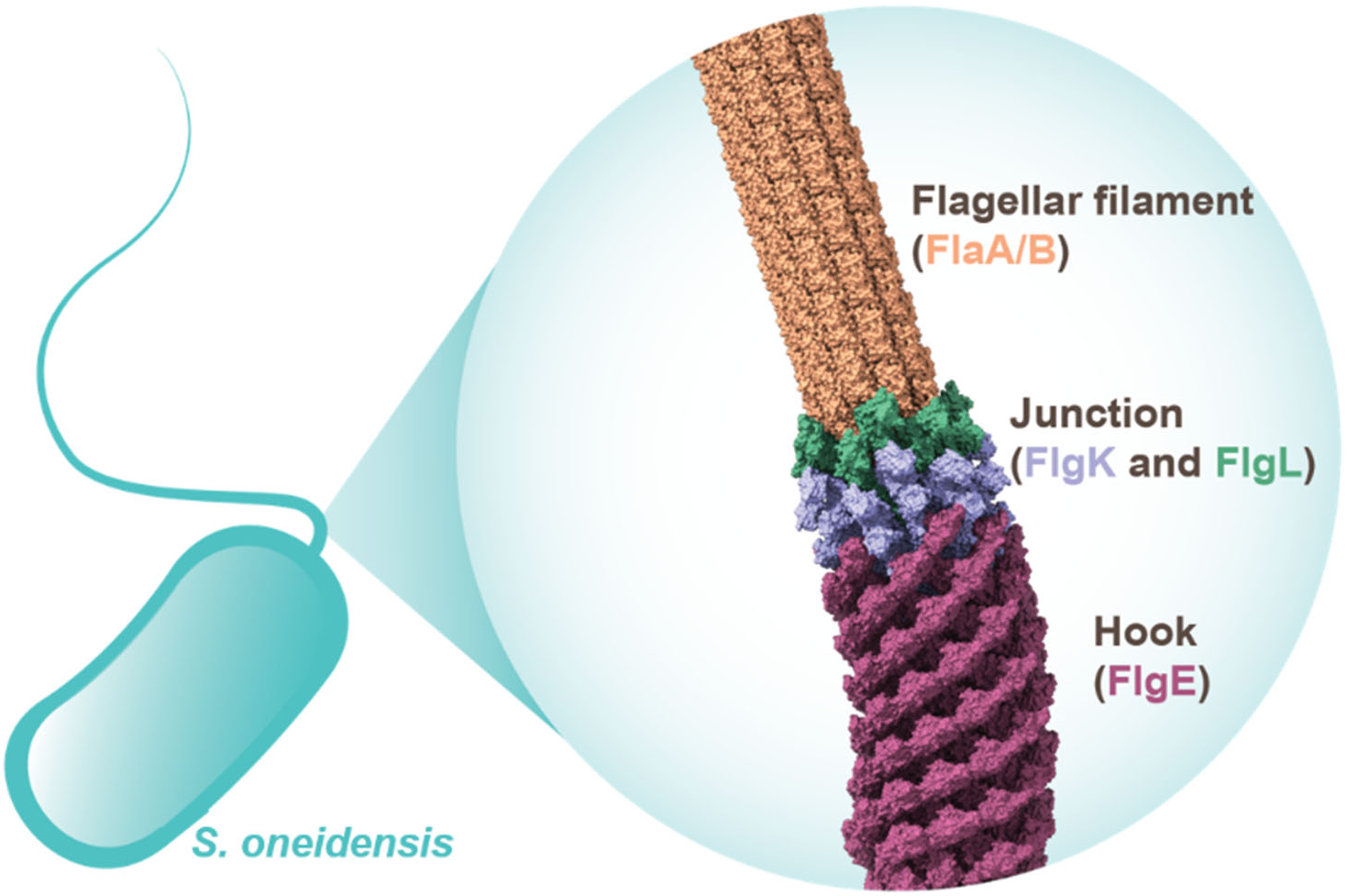

## Introduction

Motility is often vital to a bacterium’s survival in diverse environments. In liquid media, bacterial motility is driven by extracellular flagella^1–3^, whose corresponding propulsion strategy has inspired the engineering of motile microbots^4, 5^. The extracellular bacterial flagellum comprises the following components: the basal body, hook, hook-filament junction (HFJ), filament and cap^6^. Among these, the basal body, hook, HFJ and filament are primarily responsible for bacterial motility, listed in order of energy generation and transfer^7^. The basal body contains a rotor-stator complex that harnesses the ion gradient across the cytoplasmic membrane to generate mechanical energy, powering the propeller-like motion of the flagellum^8^. Connected to the basal body is the hook, a curved segment composed of multiple copies of the hook protein FlgE^9^. The hook directly connects to the rod in basal body and transmits the torque from the basal body to the rest of flagellum^9^. The structures of non-flagellated hook in *Campylobacter jejuni* and *Salmonella enterica* have been determined by cryogenic electron microscopy (cryoEM)^10–12^. Attached to the distal end of the hook is the HFJ consisting of hook-associated proteins FlgK and FlgL that together function as gaskets between hook and filament^7^, which ensures proper mechanical linkage. The structures of hook-associated proteins in *X. campestrisI*^13^, *L. pneumophila*^14^, *B. pseudomallei*^15^, *C. jejuni*^16, 17^ and *S. enterica*^17^ have been solved by X-ray crystallography or cryoEM. Further along, the filament itself is a helical assembly that propels the bacterium. Due to technical challenges in resolving cryoEM structures of curved filaments, structural studies of the flagellar filament have mostly been limited to straight-filament mutants that comply to strict helix symmetry^18–20^.

Movement of *Shewanella oneidensis* MR-1 (*S. oneidensis*) in their habitats^21–23^ is driven by a single polar flagellum^24, 25^. *S. oneidensis* is known for its versatile electron-generation capability, enabling the decomposition of organics and metals. This versatility has made *S. oneidensis* a focal point in studies on bioremediation of heavy metal wastewater^26^, biosynthesis of nanoparticles with heavy metals^24, 27^ and bio-electrochemical systems^24, 27^. All these applications will potentially benefit from enhancing the motility of this bacterium that can facilitate faster attachment to metallic or organic targets. Therefore, a deeper understanding of the motility mechanism in *S. oneidensis* is prospective. Despite its significance, the structures of motility-related components in the extracellular flagella of *S. oneidensis* have not yet been resolved. Notably, *S. oneidensis*, as a multi-flagellin species, contains two types of flagellins, FlaA and FlaB^21, 28^. The structure-based functional differences between these flagellins that distinguish their roles in bacterial motility are poorly understood. In fact, it’s not rare for flagellated bacterial species (approximate 45%) to possess more than one flagellin gene^29^, but the implications of their presence within the flagellar filament remain mostly unclear. Previous structural studies on multi-flagellin species was mainly limited to helical reconstructions of engineered mutants where other flagellins are knocked-out^18, 19^. Structures involving multiple flagellins have recently been determined in *Escherichia coli* O157:H7, *E. coli* O127:H6, *Sinorhizobium Meliloti* and *Achromobacter* and show deviations from each other mainly in their outer domains^30^.

Here, we have captured in situ flagellum attached to a bacterium of *S. oneidensis* by cellular cryoET technique and report the cryoEM structures of motility-related components in extracellular flagella: the filament, hook and HFJ at resolutions ranging from 2.8 to 6.1 Å. The filament and hook consist of multiple copies of flagellins and flgE, respectively, each adopting 11 different conformers responsible for their curvature generation. The HFJ structure is composed of 11 FlgK and 11 FlgL which form mechanical linkage that reconcile incompatibility between the filament and hook. Regarding the filament, we also identify the FlaA and FlaB isoforms with their spatial distribution and we confirm the critical residues 129 and 134 differentiate motile functions of the two isoforms. These structural insights not only deepen our comprehension of the motility mechanisms in *S. oneidensis*, but also potentially benefit the application of this bacterium in bioremediation, biosynthesis, bio-electrochemical systems and inspire the design of motile microbots in medical fields.

## Results

### Overall architecture of the extracellular flagellum in *S. oneidensis*

The cryoEM image of frozen-hydrated *S. oneidensis* (**Figure 1a**) reveals a capsule-shaped bacterial cell with a single extracellular flagellum projecting from one pole. The slice and segmentation of tomogram in **Figure 1b-c** highlight the three motility-related components of an extracellular flagellum: the hook, HFJ and flagellar filament extending outward from the basal body with decreasing outer diameters. Notably, the hook is more curved than the flagellar filament, and the HFJ serves as a mechanical linkage between them. The segmentation result (**Figure 1c**) shows that the flagellar filament forms a left-handed supercoil throughout the extracellular flagellum of wild-type *S. oneidensis*.

**Figure 1.**
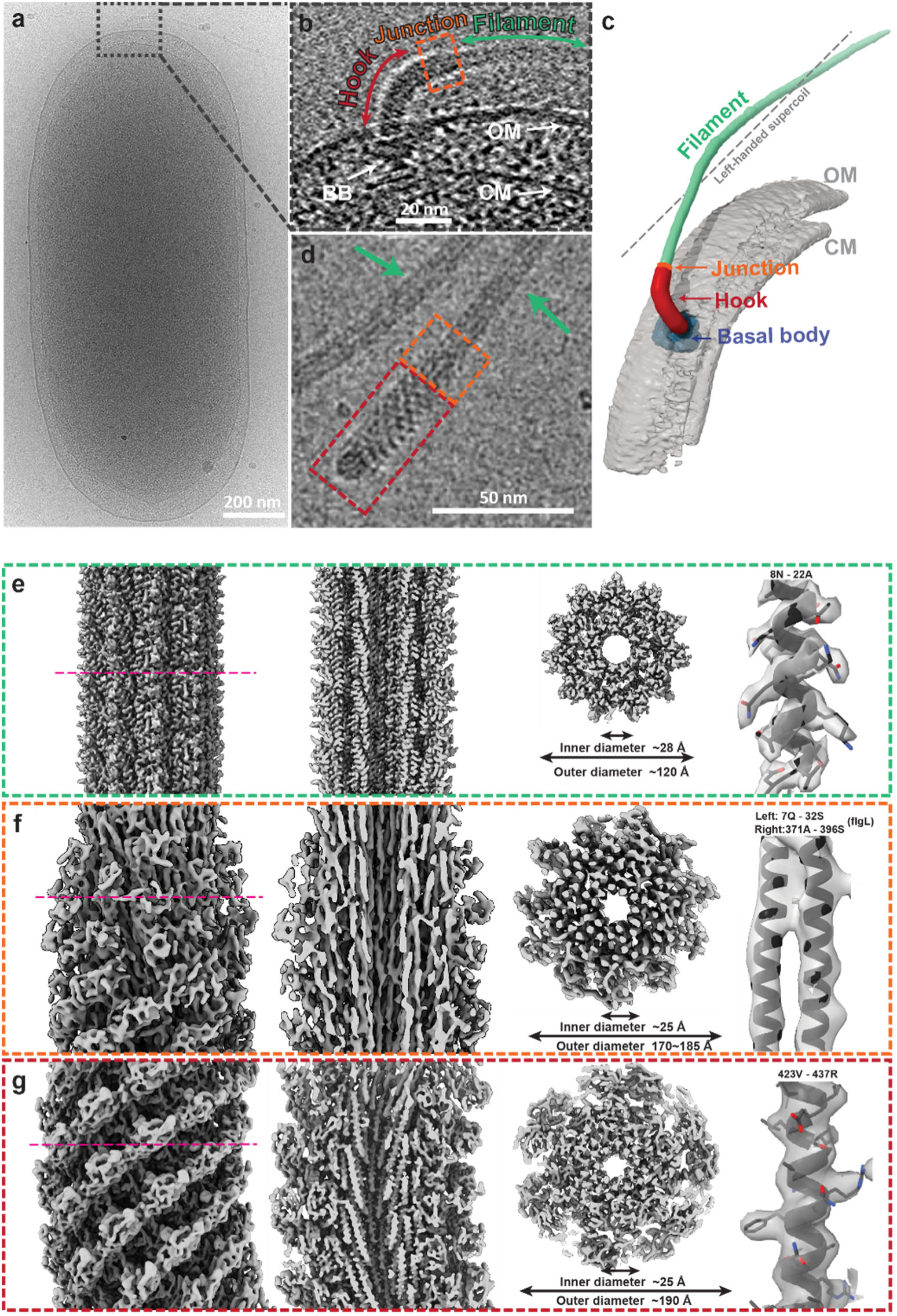
Overall structure of the extracellular flagellum in *Shewanella Oneidensis.* **(a)** CryoEM micrograph containing a whole cell of *S. Oneidensis*. **(b, c)** Tomogram slice **(b)** and segmented map **(c)** illustrating intact hook (FlgE), HFJ (FlgK and FlgL) and filament (FlaB and FlaA). **(d)** Representative cryoEM micrograph of purified extracellular flagella, highlighting hook (red square), HFJ (orange square) and filament (green arrows). **(e-g)** CryoEM maps of flagellar filament **(e)**, HFJ **(f)** and hook **(g)**. Showing from left to right: side view, axial cross-section, radial slab and regions of representative density with fitted models. Radial slab sectioning surfaces are indicated by magenta dashed lines in the side views.

To investigate the molecular basis of flagella-driven motility in *S. oneidensis*, we purified the extracellular components of the flagella (**Figure 1d**) and obtained their corresponding high-resolution cryoEM structures by single-particle analysis (SPA) (**Figure 1e-g**). With helical symmetry reconstruction, we obtained a 2.8 Å resolution density map for the flagellar filament, which clearly resolves amino-acid side chains (**Figure 1e**). Additionally, we reconstructed the asymmetric structures of HFJ and hook at resolutions of 6.1 Å and 4.2 Å, respectively (**Figure 1f-g**). All these structures feature an axial tunnel with an inner diameter of 25-28 Å, while their outer diameters range from 120 to 190 Å (**Figure 1e-g**). These structural insights have allowed us to uncover the molecular interactions among their subunits and a mechanistic understanding of *S. oneidensis* motility, as detailed below.

### Identification of FlaA and FlaB isoforms in native flagellar filament

Previous proteomic studies in *S. oneidensis*^21^ show that the flagellar filament is a polymer of two different flagellins: FlaA and FlaB, with 89% sequence identity (**Figure 2a**). Using a similar helical cryoEM reconstruction strategy of other bacterial flagellar filaments^18, 19, 31^, we obtained a high-resolution structure (2.8 Å, **Figure 2b**). The filament’s helical parameters are 65.41° twist and 4.85 Å rise (**Figure 2b**). This density map resolved long side-chain densities that only exist in FlaB, such as N147, K188 and Q243 (**Figure 2c** and **S2a-b**), without detectable FlaA features. Since the overall filament is expected to be assembled with both FlaA and FlaB^21, 28, 32^. The results with obvious FlaB features in our helix reconstruction suggest that the filament predominantly contains FlaB^21^.

**Figure 2.**
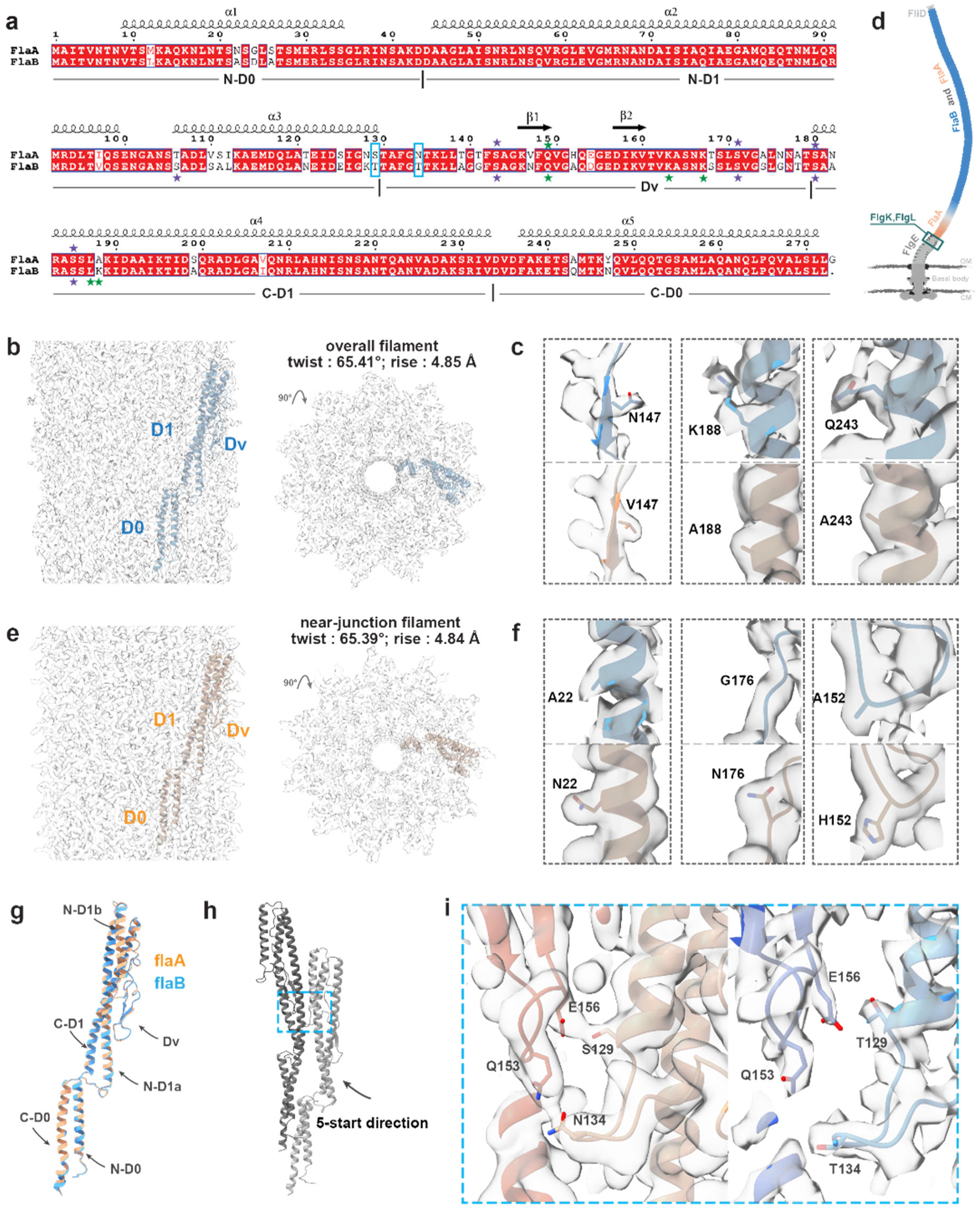
FlaA and FlaB isoforms in native flagellar filaments. **(a)** Sequence alignments of FlaA and FlaB isoforms in *S. oneidensis* (89% identity). Previously reported glycosylation and methylation sites ^33^ are marked with purple and green stars respectively; the blue squares denote residues involved in the interaction interfaces (−5 direction) in the filaments. **(b)** CryoEM map reconstructed from the overall filament with helical symmetry, with one flagellin (blue) fitted. **(c, f)** FlaA (orange) and FlaB (blue) models with density maps of representative residue 147, 188, 243 **(c)** and residue 22, 176 and 152 **(f)**. **(d)** Schematics of the distribution of FlaA and FlaB in flagellar filament based on a previous study in *Shewanella* ^29^. The orange filament segment is termed as the near-junction filament in our study. Relative colors, brightness levels and lengths in this illustration are for visual clarity and do not reflect accurate proportions or measurements. **(e)** CryoEM map reconstructed from the near-junction filament with helical symmetry, with one flagellin (orange) fitted. **(g)** The structural alignment of FlaA and FlaB subunits with 0.4 Å RMSD. **(h)** Interaction interfaces (blue dash square region) between two adjacent subunits in the 5-start directions. **(i)** Zoomed-in region from **(h)** showing that residue 129 and 134 involve 5-start interactions for the FlaA filament (orange), in contrast to no interactions for the FlaB filament (blue).

Therefore, to localize the minor flagellin component FlaA within the filament, we next focused on resolving the structure of the filament segment in the near-junction region (**Figure 2d**, orange filament segment), where FlaA is most likely to be detected based on previous fluorescent micrographs of *Shewanella*^29^. Despite a limited number of extracted particles in the near-junction region (4.6k, see Methods for details), we also applied helical symmetry and obtained a high-resolution structure (3.5 Å) (**Figure 2e**). The helical parameters of near-junction filament segment are 65.39° twist and 4.84 Å rise, which is non-identical from the overall filament (65.41° twist and 4.85 Å rise). The obtained cryoEM map reveals obvious densities of long side chains that only exist in FlaA, such as N22, H152 and N176 (**Figure 2f** and **S2a-b**), without noticeable FlaB features detected, which confirms that the filament segment in the near-junction region consists of dominant FlaA.

Such spatial arrangement of two flagellins is attributed to regulation of flagellum assembly by the timing of flagellin expression: FlaB expression is induced by the transcription factor σ^28^ (FliA) only after the assembly of hook-associated proteins (FlgL, FlgK and FliD)^32^. After the assembly of the hook-associated proteins, the flagellum initiates the growth of the filament. At this stage, FlaA subunits are available to be exported from the basal body, while FlaB expression has just begun^29, 32^. Consequently, FlaA is concentrated in the near-junction region, and the helix reconstructed density map represents FlaA filament structure (**Figure 2d**) termed as the near-junction filament in our study. And since the whole filament consists of dominant FlaB, its corresponding helix-reconstructed map represents the structure of FlaB filament. By comparing the two resolved structures, we could investigate the structural differences between FlaA and FlaB.

With an RMSD of just 0.4 Å, the monomeric structures of FlaA and FlaB are highly conserved (**Figure 2g**), in contrast to significant structural deviation of flagellins in other multi-flagellin species^30^. Both FlaA and FlaB comprise only two domains: D0, located closer to the central tunnel, and D1, forming the outer surface (**Figure S1a**). As shown in **Figures 2g** and **S1b**, each monomer contains five α-helices: N-D0, N-D1a, N-D1b, C-D1 and C-D0, connected by loops. The region between N-D1b and C-D1 is named as the hypervariable domain (Dv), containing β-hairpin secondary structure elements^21^.

Inter-subunit interactions in the FlaB filament are shown in **Figure S1c-g**; similar interactions also exist at the same locations in the FlaA filament. However, in FlaA filament, there are newly identified inter-domain interactions between Dv regions in the 5-start direction (**Figure 2a, h-i**), involving **S129**-E156 and **N134**-Q153. Such interactions are absent in the FlaB filament due to different residues at 129 and 134 (**Figure 2i**). These two critical residues (highlighted in blue squares of **Figure 2a**) involved in this interface (Dv regions in the 5-start direction), predominantly account for the functional differences in motility between FlaA and FlaB in *S. oneidensis*^21^. Mutations swapping these residues demonstrate that FlaB^T129S^ or FlaB^T134N^ decreases motility in the FlaB-only strain, while FlaA^S129T^ or FlaA^N134T^ enhances motility in the FlaA-only strain^21^. The above four cases exhibit similar motility levels, suggesting that residues 129 and 134 are the major determinants of motility differences between FlaA and FlaB^21^. Such motility distinctions primarily originate from additional interactions in FlaA filament between neighboring Dv domains involving residues 129 and 134.

Moreover, certain post-translational modifications differ between FlaA and FlaB. Extra non-protein densities corresponding to reported methylation sites were found only in FlaB at K167 and K189 (**Figure S2c-d**)^28, 33^. Densities related to reported O-link glycosylation sites^33^ are identified in both FlaA and FlaB at residues T/S106, S143, S173, S180 and S185 (colored magenta in **Figure S1b, Figure S2e-f**). However, according to previous mass spectrometry data^28^, glycosylation at residue 106 was detected only in FlaB, possibly due to the lower abundance of FlaA in overall filament. Neither glycosylation nor methylation was reported to significantly affect bacterial motility^33^, which might play other roles, such as protection imparted by glycosylation.

### Curvature generation and supercoiling of flagellar filament

Filament supercoiling determines the wave function of flagellar propelling and is thus central to bacterial motility. The curvature of our helix-reconstructed flagellar filament was lost (**Figure S1a**) due to imposition of helix symmetry^34^. Therefore, we reconstructed an asymmetric (i.e., C1) structure at 3.2 Å resolution (**Figure 3a**, left) following a data processing strategy similar to that described previously^34^. This map exhibits a slight curvature (**Figure 3a**). Since our study on curvature generation focuses on the conformational deviations of the Cα main chains of flagellins, we constructed a corresponding Cα model by disregarding the side-chain differences between the two flagellin isoforms. We then assembled the resolved curved segment models to reconstruct complete helical turns of the filament, forming a left-handed supercoil with helical radius (0.2 µm) and pitch (1.7 µm) (**Figure S3a**).

**Figure 3.**
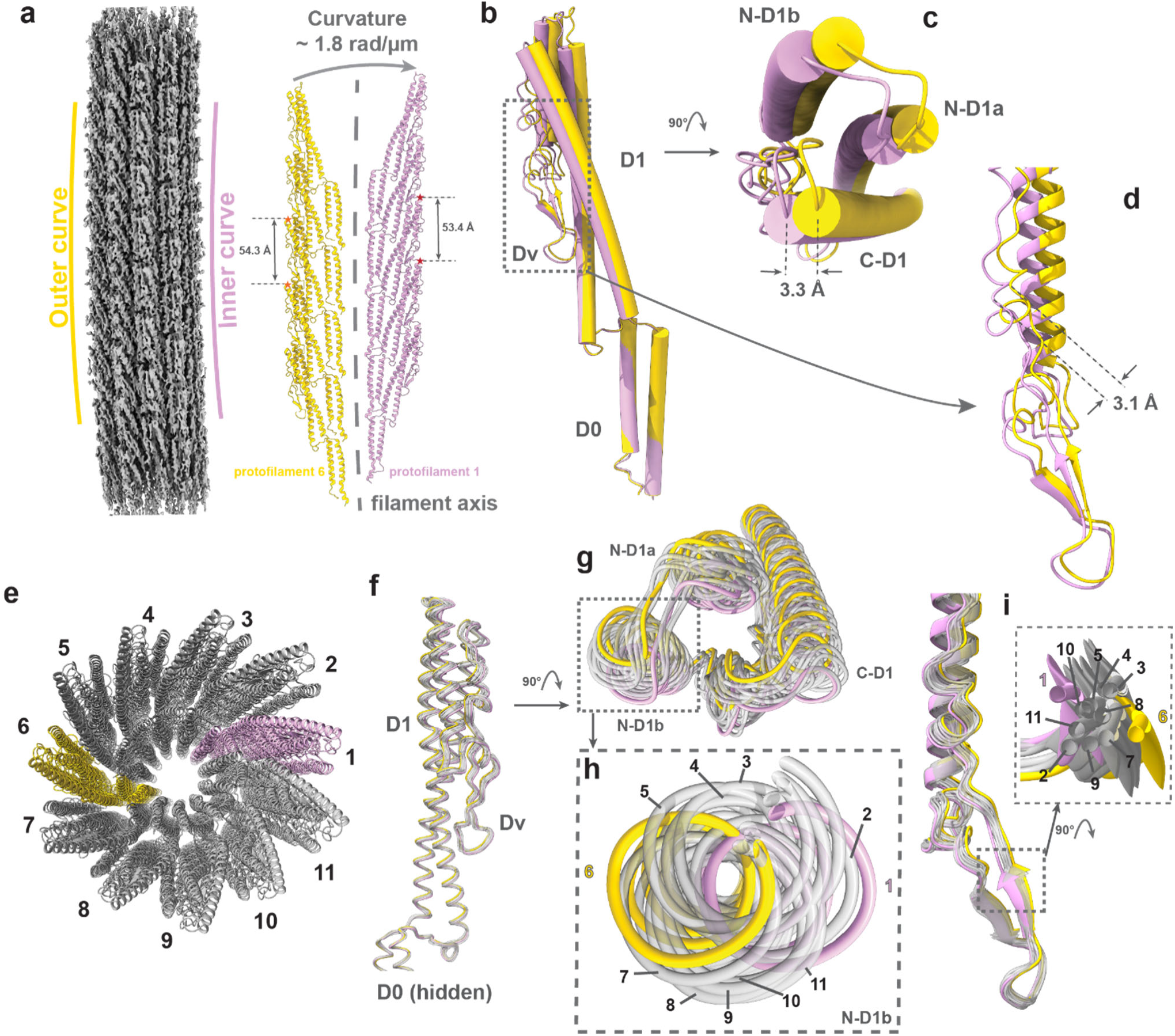
Curvature generation of flagellar filaments. **(a)** CryoEM density map of the curved flagellar filament, and models of the outermost (yellow) and innermost (pink) protofilaments. Stars mark residue 180 in the subunits. **(b)** Alignment of the outer (yellow) and inner (pink) subunits by D0 domain. **(c)** Top view of the D1 domain from **(b)**. **(d)** Zoomed-in view of the Dv domain from **(b)**. **(e)** Top view of the filament, showing its 11 protofilaments with the outermost flagellins in yellow (#6), the innermost flagellins in pink (#1), and the rest in grey. **(f)** Alignment of subunits in 11 flagellins by D0 domain. **(g)** Top view of D1 domains rotated 90° from **(f)**. **(h)** Zoom-in view of nearly circular pattern from **(g)**. **(i)** Zoom-in view of the Dv domains from **(f)**. The inset shows the deviation of 11 conformers.

The model shows a curvature, which forms the supercoiling of overall filament, of about 1.8 rad/ µm. The model bends from outermost (yellow) towards innermost (pink) protofilaments (defined as the flagellins stacking along the filament axis^35^, **Figure 3a**), which is caused by two structural factors. Firstly, the inter-subunit distance between neighboring subunits within each protofilament is different, inferring that protofilaments exhibit the packing difference of subunits (**Figure 3a**, representative distance labeled between D1 domains). Besides, the intra-subunit flexibility of protofilaments also contributes to the curvature. The different conformations can be visualized when the two corresponding subunits—one from the outer (yellow) and other from the inner (pink) protofilament—are aligned by their D0 domains (**Figure 3b**). Both the D1 (**Figure 3c**) and Dv (**Figure 3d**) domains in the innermost subunit extend farther away from the filament axis compared to the corresponding domains of the outermost subunit. The conformational deviation is becoming more noticeable in the area that is farther away from the filament axis of the flagellar filament, such as the selected farthest positions in Dv with the largest shift of 3.1 Å and in D1 with the largest shift of 3.3 Å (**Figure 3c-d**).

Our structure shows that the subunits along one protofilament share nearly identical conformation, as also demonstrated for *E.coli* K-12 flagellar filament^34^. Therefore, we only compared the representative conformers from all the 11 protofilaments of our model (**Figure 3e**; subunits in pink and yellow are the innermost and outermost subunits, respectively) by aligning their D0 domains of the representative subunits from these protofilaments (**Figure 3f**). As shown in **Figure 3g-h**, the 11 aligned conformers exhibit a nearly circular pattern, suggesting that the subunit undergoes a highly constrained hinge-type of transformation pivoting around D0. As for the hypervariable domain (Dv), the transformation is not in a clear trajectory as the D1 domain (**Figure 3h, i**).

The packing difference across each protofilament, together with the conformational deviation of subunits, results in the curvature with 11 conformers from different protofilaments. Meanwhile, the near-identical interactions between neighboring subunits along the same protofilament allow synchronized supercoiling throughout the whole overall filament.

### Curvature of the hook — a biological universal joint

Previous hook structures from multi-flagella bacteria are inferred from non-flagellated polyhooks^10–12^; the 4.2 Å resolution density map of the hook segment in our current work offers an opportunity to understand subunit arrangement and supercoiling mechanism for the wild-type hook in the single flagellum of *S. oneidensis*. We built the atomic model of the hook segment by rigid-body fitting the individual domains from AlphaFold2-predicted FlgE model into the density map, and then refined the composite model with connected domains using *Phenix*^36^.

To gain insights into the supercoiling of the hook, we constructed the supercoil model by stacking up the resolved structure of hook segment (**Figure S3b**). The helical radius and pitch of this left-handed supercoil are around 26 nm and 139 nm, respectively. In contrast to the filament length in micrometer scale, the length of hook is estimated based on our tomogram to be about 55 nm (**Figure 1c**), which is consistent with that of *Salmonella*^37^. If the hook length is far below 55 nm (≤42 nm), it would be too stiff to function as universal joint. By contrast, an increased length (≥75 nm) would destabilize the hook and therefore increase the probability of its unnecessary buckling^38^. Therefore, the length of hook is an optimum solution to facilitate efficient torque transmission in their native environments, such as aquatic and sedimental environments.

Hook is composed of multiple copies of FlgE, which contains D0, D1 and D2 domains with an additional Dc domain^12^ (**Figure 4a**). Recall that we found above that the structures of flagellins within the same protofilament are nearly identical but those in different protofilaments differ. We suspect that these structural differences of subunits contribute to the curvature of the filament. Accordingly, we first analyzed the packing difference among subunits in our model. The model is colored with respect to the Cα distance between corresponding residues of two adjacent subunits along protofilament (**Figure 4b**), the subunits along the same protofilament share nearly identical distance. The subunits are more closely stacked in the innermost protofilament (IMP, compressed form) than those in the outermost protofilament (OMP, extended form), which results in the bending towards the IMP. Such compression and extension of protofilaments are associated with 11 FlgE conformers residing in 11 protofilaments (**Figure 4b**). The conformational deviation among the subunits along the same protofilament is negligible which is consistent with *Salmonella*^12^. To explore the origin of the conformational deviation, we compared innermost #1, intermediate #4 and outermost #7 conformers by superposing their D0 domains (**Figure 4c**) and indicate that the loops connecting three domains act as two flexible hinges (in black dashed squares) and enable conformational changes of the subunits. By comparing representative IMP and OMP subunits, the tilting range of FlgE is up to 14° by taking hinge 1 as origin, and the shift increases from D1 (5 Å) to D2 (25 Å) domains (**Figure 4d**), in contrast to the constrained filament conformers (up to 3.3 Å shift). Thus, the relatively rigid domains connected with flexible hinges of FlgE result in 11 conformations from 11 protofilaments, which cause the compression or extension of protofilaments.

**Figure 4.**
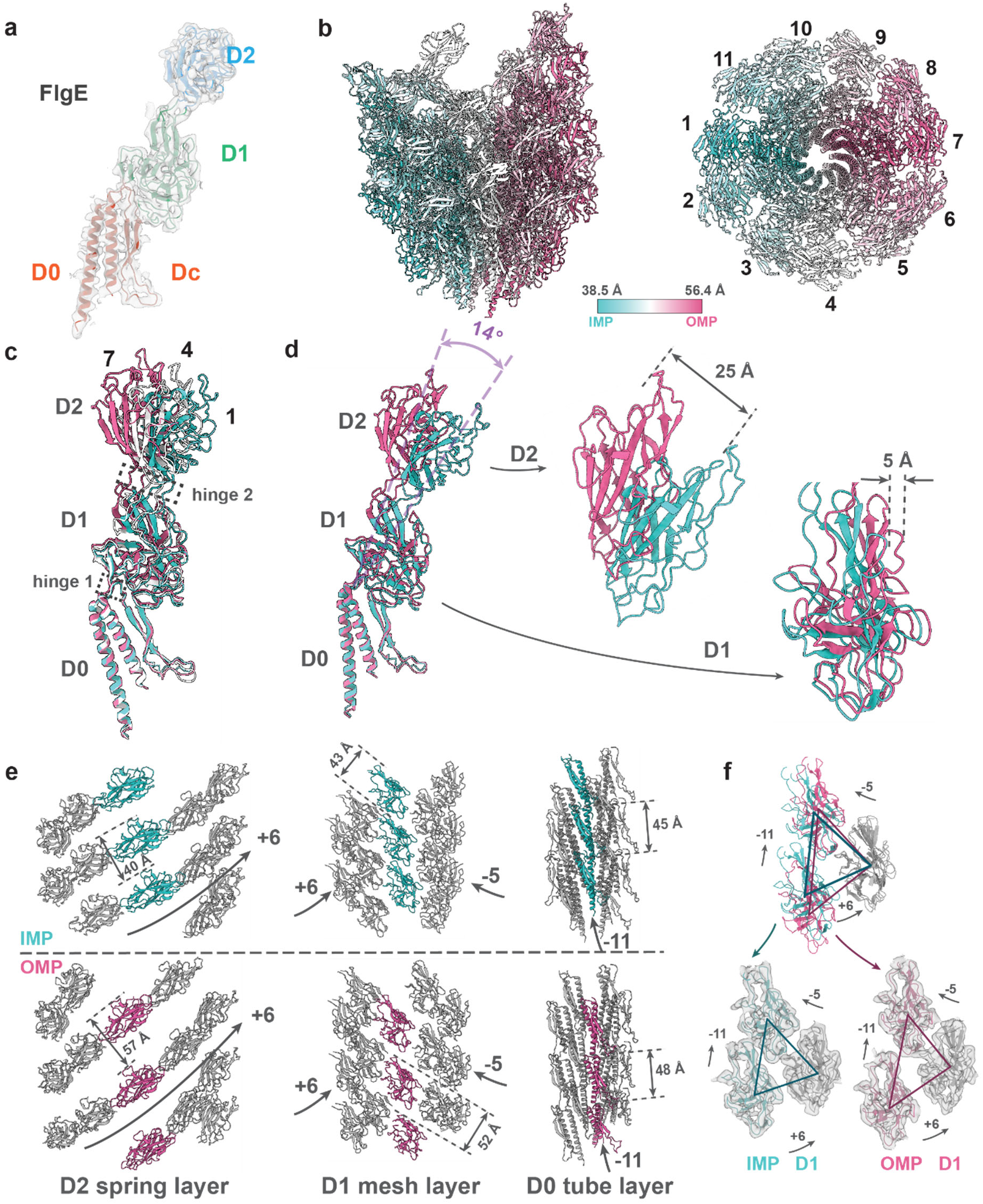
Supercoiling of wild-type hook. **(a)** FlgE monomer colored by different domains fitted in density map. **(b)** Side and top views of hook. The structures are color-coded regarding the distance between residue 249 of two adjacent subunits along protofilament. Cα distances of IMP (compressed form) and OMP (extended form) are 38.5 Å and 56.4Å respectively. **(c)** Alignment of subunits in IMP (#1), OMP (#7) and intermediate state (#4) by their D0 domains. **(d)** Alignment of subunits in IMP (#1) and OMP (#7) by D0 domains. And zoom-in views of the deviations in D1 and D2 domains. **(e)** Three layers show the arrangements of D2, D1 and D0 domains from IMP and OMP sides. From left to right: D2, D1, D0. **(f)** Structural comparison of the three nearest D1 domains near IMP (cyan) and OMP (magenta) aligned by the D1 domains (grey).

To better elucidate the role of each FlgE domain in hook as a universal joint which simultaneously exhibits bending flexibility (degree to bending deformation) and twisting rigidity (resistance to torsional deformation), we then analyze the arrangements and associated interactions of domains layer by layer across domains D2, D1, D0, as detailed below (**Figure 4e, S4a-b**).

As for the outermost domains D2, the variable spacing between them along the same protofilament, measuring from 40 Å in IMP to 57 Å in OMP (**Figure 4e**, left column), allows for the high bending flexibility of hook. D2 domains tightly connect to the adjacent D2 in the 6-start direction while no interaction is observed in other directions (**Figure S4a**, left column). Residues potentially mediating interactions between D2 domains are identified only in 6-start direction (**Figure S4b**, left column) based on reasonable Cα atom distances. However, the involved residues vary across protofilaments, such as N219 - A304 in IMP shifted to S275 - A304 in OMP (**Figure S4b**, left column). With D2 domains tightly connected in 6-start direction but loosely packed in others, they form a spring-like arrangement in the outermost shell of the hook (**Figure 4e**, left column), similar to that observed in *Salmonella*^11^. This D2 spring layer is also indispensable for the curvature of hook, as demonstrated by a mutation experiment in *Salmonella* where the deletion of D2 resulted in a straight hook^39^.

Regarding the middle layer composed of D1 domains, the spacing deviation between D1 domains along protofilaments is less than that of D2, reflected by the distances from 43 Å in IMP to 52 Å in OMP compared to that from 40 Å to 57 Å between D2s (middle column, **Figure 4e**). Thus, the D1 domains form a less flexible layer than D2. To further analyze the arrangements of D1 domains, the structural comparison of three selected D1 domains near the IMP and OMP are shown in **Figure 4f**. The D1s on the top and lower-right are at comparably stable positions, maintained by the constitutive interactions between D1s in the 5-start direction (**Figure S4b**, middle column). However, the relative flexible positions of D1s in 6-start and 11-start directions (**Figure 4f**) resulted from the lack of interaction in 6-start direction and the dynamic interactions at the D1-D2/D1 interfaces along 11-start direction (**Figure S4c-d**), Therefore, D1 mesh layer stabilizes the hook by maintaining the connections in 5-start direction while preserving a certain degree of flexibility in other directions.

The innermost D0 layer exhibits the most rigid structure across three radial layers, minimizing the difference of adjacent subunit distances in IMP (45 Å) and OMP (48 Å) along protofilaments (**Figure 4e**). Unlike the other two layers, interactions between D0s occur along 11-start direction dominated by stable hydrophobic interactions (**Figure S4b**, right panels).

Besides, the Dc region (D0: 27-79 residues) persistently interacts with D1 in 5-start direction (**Figure S4e-f**). Therefore, with the longitudinal D0-D0 linkages and inter-layer Dc-D1 connections, the D0 tube layer serves as a rigid scaffold, primarily contributing to the stability and twisting rigidity of hook.

The distinct domain lattice within each layer (**Figure 4e**), together with inter-layer and intra-layer interactions (**Figure S4**) that integrate both structural stability and dynamic adaptability, collectively facilitate the synchronized conformational changes of subunits, enabling the coordinated compression and extension of the 11 protofilaments during rotation. Thus, the hook, as a biological universal joint with high bending flexibility and twisting rigidity, transmits rotational torque efficiently from flagellar rod to the propelling components in extracellular flagellum.

### Molecular composition and structural arrangement of the hook-filament junction complex

The HFJ proteins: FlgL (**Figure 5a**) and FlgK (**Figure 5b**), connect the flagellar filament and hook, and are the prerequisites for the formation of a motile flagellum^13, 17^. We dock individual domains from AlphaFold2-predicted models into the map (6.1 Å) with rigid-body fitting. 11 FlgL and 11 FlgK together constitute the HFJ (**Figure S5a-b**), consistent with previous stoichiometry analyses in *Salmonella* and *Campylobacter*^17, 40^.

**Figure 5.**
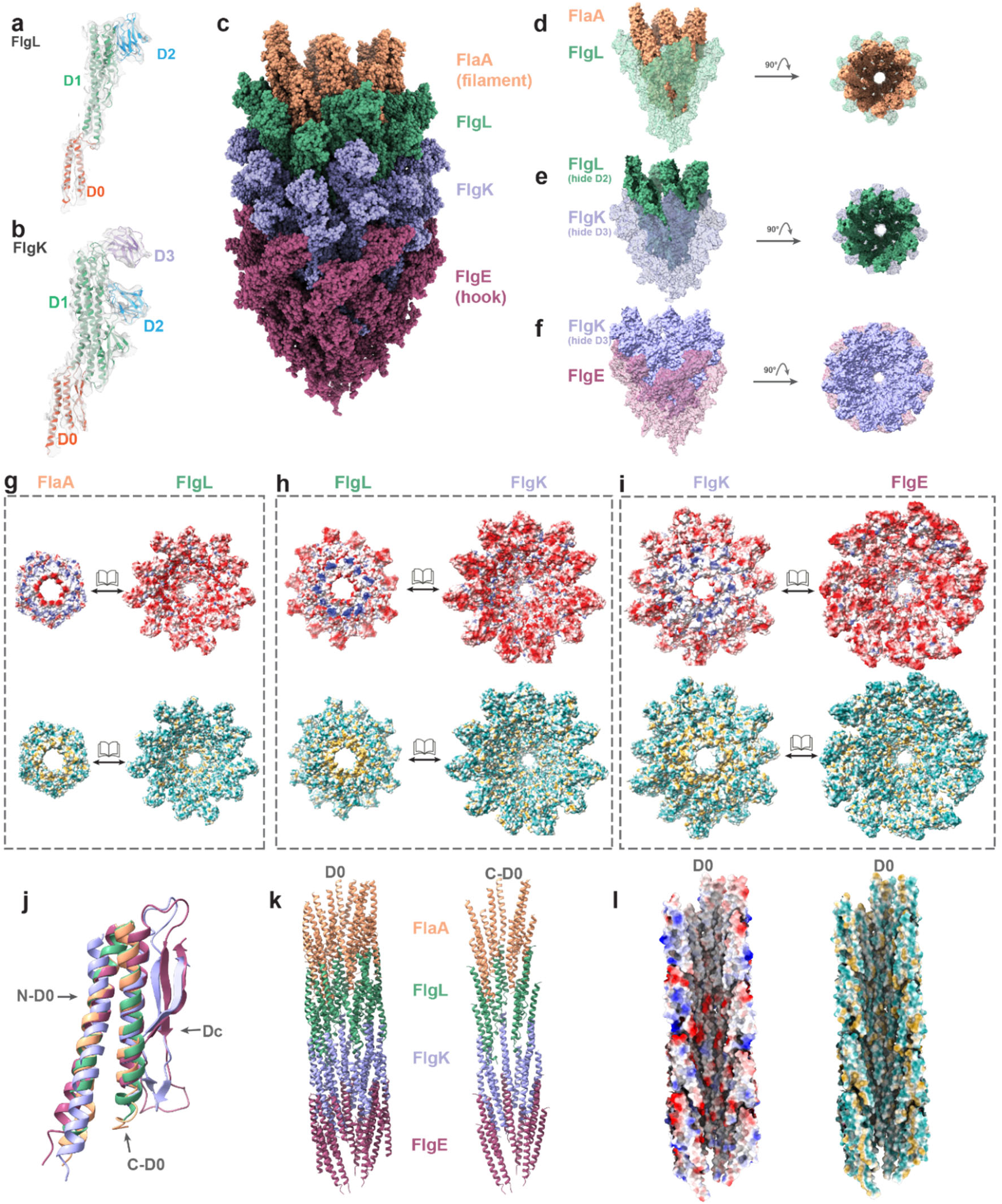
Native structure of hook-filament junction complex. **(a)** FlgL model colored by different domains fitted in density map. **(b)** FlgK model colored by different domains fitted in density map. **(c)** Modelling of hook-filament junction complex, including one helical turn each of FlaA, FlgL, FlgK and FlgE. **(d-f)** The models of FlaA - FlgL **(d)**, FlgL - FlgK **(e)** and FlgK – FlgE **(f)** helical turns in nest form. **(g-i)** The interfaces of FlaA - FlgL **(g)**, FlgL - FlgK **(h)** and FlgK - FlgE **(i)**, shown in the form of electrostaticity and hydrophobicity patterns. **(j)** Alignment of D0 domains across FlaA, FlgL, FlgK and FlgE. **(k)** Structurally conserved D0 tunnel in the HFJ region, with C-D0 domains forming the inner tunnel surface. **(l)** The electrostaticity and hydrophobicity of D0 tunnel, showing the properties of inner tunnel surface.

The 250Å-long HFJ consists of two gasket-like structures connecting the filament and hook (**Figure 5c** and **S5c**). The FlgL gasket interfaces with flagellins (FlaA) of the filament, and the FlgK gasket with the hook proteins (FlgE) (**Figure 5c** and **S5c**). The outer diameter of the FlgL gasket is ∼170 Å whereas that of the FlgK gasket is around 185 Å (**Figure S5a-b**). These diameters fall within the range between the hook (∼190 Å) and the filament (∼120 Å). The four ring-like structures, composed of FlaA, FlgL, FlgK and FlgE respectively, are nested within one another (**Figure 5d-f**), forming a smooth transition from hook to filament. Except for the D2 domain in FlgL and D3 domain in FlgK, all other domains are engaged in linkages throughout four rings (**Figure 5d-f**), which is supported by the HFJ in *Salmonella*^17^ that lacks the D2 domain in FlgL and D3 domain in FlgK. All interfaces between four ring-like structures display potential electrostatic interactions mainly in outer domains (D1 and D2), and prominent hydrophobic interactions in the core region (D0) (**Figure 5g-i**).

The D0 domains of these four proteins all contain two α helices. Additionally, FlgK and FlgE have an extra Dc region that consists of β hairpins and loops (**Figure S5d-g**). The relatively conserved structures of the two α-helices (**Figure 5j**) in D0 domains from all subunits interact through hydrophobic interactions (**Figure 5g-i**, lower panels) to form a 25-28 Å diameter central tunnel of the flagellum. The D0 domains surrounding the tunnel act as a scaffold to maintain the flagellar integrity while the torque is being transmitted from the hook to filament (**Figure 5k**, left). The C-D0 helices (**Figure 5k**, right) form the inner surface of the tunnel. Notably, polar residues lining along the inner surface of this narrow tunnel (**Figure 5l**) would allow partially unfolded flagellins to pass through without aggregation or clogging, facilitating efficient export of flagellins for filament growth^41^.

Besides the structurally conserved D0 domains from flagellin to hook subunit in our HFJ model, the other common domain, i.e. D1, also exhibits a transition of structural similarity (**Figure S5d-g**). These structural insights further confirm previous speculation that the linkage between four types of proteins in HFJ complex is mediated by quasi-homotypic interactions that mimic the homotypic flagellin-flagellin and FlgE-FlgE interactions^13^. Therefore, FlgL and FlgK gaskets act as a transition zone to reconcile the incompatibility between the hook and filament.

## Discussion

Our structures of the extracellular components of *S. oneidensis* flagellum (**Figure 6a**) provide insight into the motility mechanism of monotrichous bacteria through helical waveforms. The coordinated actions of the hook, filament and HFJ of the single extracellular flagellum impart both bending flexibility to facilitate propelling motion and structural integrity to resist twisting deformation. The whole extracellular flagellum is composed of 11 protofilaments that differ subtly in the levels of either compression or extension. Such compression and extension give rise to microscopic curvature and macroscopic supercoiling, which are key to motility. Besides, the D0 domains of all three components form a tubular backbone (**Figure 5k** and **6b**) via mainly hydrophobic interactions along the protofilament (**Figure 5g-i**, lower panels), which can resist extreme bending and twisting during propulsion to prevent deformation.

**Figure 6.**
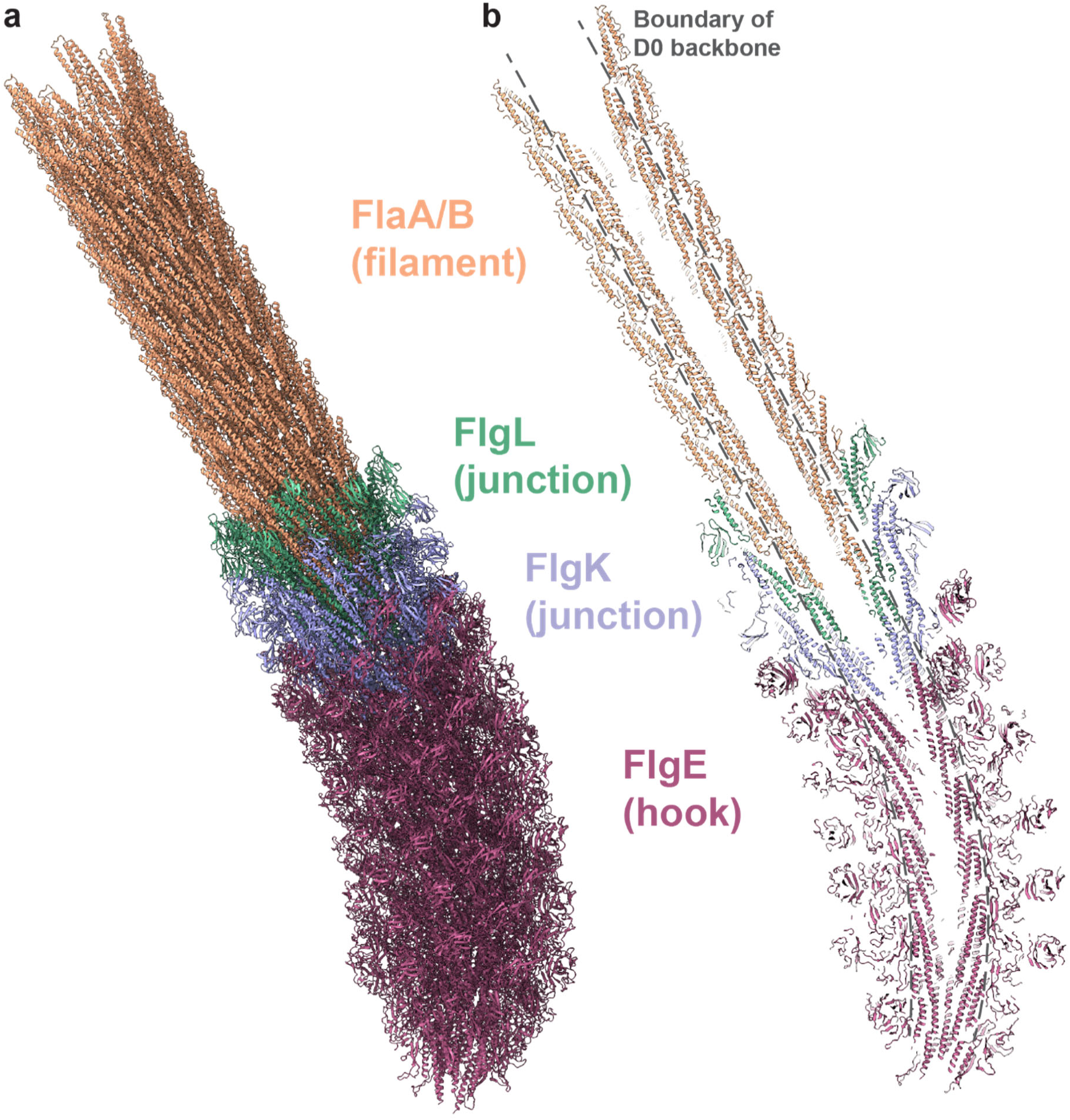
Modelling of the extracellular flagellum. **(a)** Pseudo-atomic model of stacked extracellular flagellum, containing hook (curvature ∼22 rad/µm), HFJ and filament (curvature ∼1.8 rad/µm). **(b)** A section of the model, showing the central tunnel and domains. The boundary of D0 backbone is lined out with grey dash lines.

Each of the three components plays a distinct and essential role in motility. The hook, with domains of its subunits obviously stacked loosely [especially, D2 are arranged like springs and D1 are networked as a mesh layer (**Figure 4e**, left and middle panel)], is capable of large bending. Thus, it functions as a universal joint, transferring torque efficiently from the basal body to the rest of the flagellum even though they are not coaxial. In addition, a certain level of elasticity of the hook is also crucial for turning swimming direction in monotrichous bacteria^42, 43^. By contrast, the less curved filament structure is more densely packed than the hook (**Figure 6a**), and the flagellins across different protofilaments are less flexible than the hook subunits (**Figure 3b-c** and **Figure 4c-d**). Consequently, the filament is more rigid against both bending and twisting than the hook and thus acts as the propeller. Despite the differences between the hook and filament in structures and mechanical properties, they need to function synchronously within the whole flagellum to generate propelling force through interactions to external fluids and thereby moving the bacterium forward. Therefore, HFJ with two gasket-like layers of 11 FlgK (**Figure S5f**) and FlgL (**Figure S5e**) serves as a transitional zone to counter incompatibility between the hook and filament; this interpretation is supported by our observed gradual transition of structural similarity across these four types of subunits (**Figure S5d-g**). Together, these three components coordinate to achieve propeller-like motion and facilitate bacterial motility.

Notably, *S. oneidensis* is a multi-flagellin bacterium, and we show in this study that the structures of its two flagellin isoforms are highly similar: each containing only two core domains (D0 and D1), thus differing from the situation in previously reported multi-flagellin structures of other species which typically contain more than two domains^30^. The composition and structure of the filament are influenced by bacterial habitats. According to the filament structures in our study (**Figure 2**), interactions along the 5-start direction that take place in the FlaA filament are absent in the FlaB filament, which are suggested to be the key factors influencing both overall morphology and motile function in these two types of filaments. Such difference between the two filaments aligns with flagellin mutagenesis results from another closely related species in *Shewanella*^29^ (see sequence and structural alignments between the flagellins in *Shewanella putrefaciens* and *Shewanella oneidensis* in **Figure S6**). FlaA-only filaments had smaller pitch (∼1.2 µm) and radius (∼0.17 µm) parameters than FlaB-only filaments (pitch: ∼1.79 µm and radius: ∼0.3 µm). The estimated supercoiling pitch and radius (**Figure S3a**) of our wild-type filament fall within these ranges respectively. *Shewanella* adopts different types of motion depending on the environments^22, 44^, which offers insight into the origin of its unique spatial arrangement of the flagellin isoforms. Compared to the free-swimming motion of *Shewanella* in liquid, screw-like motion, which requires wrapping the flagellum around the cell, is important for this bacterium to escape traps in structured environments like sediments^44^. Free swimming motion occurs in the wild-type, FlaA-only and FlaB-only mutants. However, the FlaA-only mutant is incapable of screw-like motion, unlike the wild type and FlaB-only mutant^29^. Therefore, the presence of near-junction FlaA segment in wild type filament helps stabilize flagellum by preventing premature screw formation during free swimming^29^. Collectively, the spatial arrangement of flagellins in the wild-type filaments in *S. oneidensis*, featuring a FlaA segment near the HFJ and dominant FlaB in the rest of the filament, defines the supercoiling and balances between swimming propulsion and screw-like motion, which enables efficient motility of *S. oneidensis* across various natural environments.

From the perspective of convergent evolution, genes of different microbial origins could evolve to adopt a convergent architectural solution to similar tasks. In the case of motility, unrelated molecules across the three domains of life polymerize into filamentous appendages to cells that seem to have converged as propulsive nano-machines to enable movement in various environments. To be specific, the major motile organelles of microbes are archaella (i.e. archaeal flagella) in Archaea, flagella in Bacteria and cilia in Eukaryotes. In Archaea, as exemplified by *Methanospirillum hungatei*, the subunit of archaellar filament deviates from that of the bacterial flagellum structurally and rather bears more similarity to the subunit of bacterial type IV pilus^45–47^. Besides rotary motion, archaellum can also enable twitching motion like bacterial type IV pili by attaching to a surface, and then followed by extending or shrinking^47^. In Eukaryotes, cilium is the filamentous motile machinery and contains a compartmentalized axoneme responsible for ciliary motion^48^. The dynein motors which power the motion are arranged along axoneme with periodicity^48^ and coordinate to produce various types of motion, such as spiral motion for *M. pusilla*^49^, in-plane bilaterally beating for *C. reinhardi*^50^, and bihelical motion for *T. brucei*^51^. Besides, the architectures of archaellum and flagellum are much simpler than that of cilium, thus are more attractive as biomimetic motility nano-machines. However, the post-translational modifications in archaella are far more diverse than in the filamentous structures in the other two domains^52^, which allow adaptation to extreme environments; such modifications introduce additional constraints and undesirable complications to the biomimetic engineering based on such machines. By contrast, the bacterial flagellum, with its streamlined structure and functionality across diverse environments, offers a more suitable template for designing versatile motile systems.

Therefore, engineering principles can be proposed based on the observations of our flagellar structures. First, bacterial flagellum achieves motility with just a few evolutionarily related types of proteinaceous subunits or building blocks. These subunits separately construct different modules for specific mechanical tasks, i.e. universal joint, junction and propeller. Those subunits may evolve from a common gene origin, and thus they retain a certain level of structural similarity to impart synchronous actions (**Figure S5d-g**). Such an evolutionary pathway offers insights into motile-machine design by simply using a foundational building block that can be modified further to embody different functions. Furthermore, the spatial arrangement of two flagellins in *S. oneidensis* identified by our cryoEM structures is a critical element impacting the mechanical property of the filament and optimizes motility by adaptively switching between free swimming and screw-like motion. Hence, *Shewanella*’s natural strategy for efficient movement across diverse environments provides inspiration for developing more adaptable motile machines, offering potential advantages over current single-flagellin filament-based designs^53^.

## Conclusion

In this study, we have determined the native structures of motility-related components in the extracellular flagellum of *S. oneidensis*, including the hook, filament and HFJ. The hook and filament consist of multiple copies of flgE and flagellins, respectively, each adopting 11 different conformations. The hook exhibits a larger curvature and mechanical flexibility compared to the filament. The resolved HFJ structure is composed of 11 FlgK and 11 FlgL which forms gasket-like layers that reconcile incompatibility between the hook and filament. Regarding the native multi-flagellin filament, we identify the FlaA and FlaB isoforms and their spatial distribution. FlaA is more abundant in the near-junction filament, while FlaB predominates in the overall filament. Moreover, we identify critical residues 129 and 134 imparting their individual motile functions of the FlaA and FlaB isoforms. Our results not only provide the first structural description of motile machinery of the model biotechnology bacterium *S. oneidensis* but also offer a deeper understanding of general motility mechanisms and valuable insights for modifying and manipulating this bacterium for applications. Such modification and maneuverability could enhance efficiency when approaching the targets for potential applications exemplified as bioremediation, biosynthesis and bio-electrochemical systems. Furthermore, the cooperative behavior of the hook, HFJ and multi-flagellin filament can also inform the design of powerful motile microbots in medical fields for applications across non-invasive microsurgery, diagnosis and therapy.

## Methods

### Cell incubation and preparation

*Shewanella oneidensis* MR-1 was first inoculated with 20 ml of LB solution. The entire flask together with the LB solution was put in a 30 °C shaker for at least 15 h (usually overnight). 200 μL of bacteria colonies were taken out and put in another flask containing 20 ml of fresh LB solution in a 30 °C shaker for about 4 hours. 5 mL of bacteria colonies were taken out and put in another flask containing 20 ml of *S. oneidensis* medium in a 30 °C shaker for 20 hours. A centrifuge at ×2500 g for 5 min was used to collect the bacteria, which were redispersed in 1X PBS buffer.

The *S. oneidensis* medium contains (per liter of deionized water): 9.07 g PIPES buffer (C_8_H_18_N_2_O_6_S_2_), 3.4 g sodium hydroxide (NaOH), 1.5 g ammonium chloride (NH_4_Cl), 0.1 g potassium chloride (KCl), 0.6 g sodium phosphate monobasic monohydrate (NaH_2_PO_4_·H_2_O), 18 mM 60% (v/v) sodium DL-lactate (C_3_H_6_O_3_) solution as the electron donor, 10 mL of 100× amino acids stock solution, and 10 mL 100× minerals stock solution. This base medium is adjusted to an initial pH of 7.2 using HCl and NaOH and then sterile filtered using 0.22 μm vacuum driven disposable bottle top filters (Millipore). Finally, before the medium is used, 0.5 mL 100 mM ferric NTA (C_6_H_6_FeNO_6_) stock solution is added per liter of medium. The 100× amino acids stock solution contains (per liter of deionized water) 2 g L-glutamic acid (C_5_H_9_NO_4_), 2 g L-arginine (C_6_H_14_N_4_O_2_), and 2 g DL-serine (C_3_H_7_NO_3_). The 100× minerals stock solution contains (per liter of deionized water): 1.5 g nitrilotriacetic acid (C_6_H_9_NO_6_), 3 g magnesium sulfate heptahydrate (MgSO_4_·7H_2_O), 0.5 g manganese sulfate monohydrate (MnSO_4_·H_2_O), 1 g sodium chloride (NaCl), 0.1 g ferrous sulfate heptahydrate (FeSO_4_·7H_2_O), 0.1 g calcium chloride dehydrate (CaCl_2_·2H_2_O), 0.1 g cobalt chloride hexahydrate (CoCl_2_·6H_2_O), 0.13 g zinc chloride (ZnCl_2_), 10 mg cupric sulfate pentahydrate (CuSO_4_·5H_2_O), 10 mg aluminum potassium disulfate dodecahydrate (KAl(SO_4_)_2_·12H_2_O), 10 mg boric acid (H_3_BO_3_), 25 mg sodium molybdate dehydrate (Na_2_MoO_4_), 24 mg nickel chloride hexahydrate (NiCl_2_·6H_2_O), and 25 mg sodium tungstate (Na_2_WO_4_). All stock solutions are sterile filtered prior to use.

### Cryogenic electron tomography of *S. oneidensis*

An aliquot of 3 µl of the bacteria-containing sample was applied onto Quantifoil holey carbon grids (200 mesh, 3/1) that were glow-discharged for 60 s at 25 mA using the PELCO easiGlow. Using a Vitrobot Mark IV (FEI/Thermo-Fisher) set to 4°C and 100% humidity, excess sample solution on the grids was blotted away by filter paper at a blot force 2 and blot time 8 s. The grids were then plunged into a pre-cooled liquid ethane/propane mixture for vitrification. The plunge-freezing conditions and cell concentration were optimized by screening with an FEI TF20 transmission electron microscope equipped with a Gatan K2 camera.

Tilt series were collected using *SerialEM*^54^ in an FEI Titan Krios transmission electron microscope (300 kV) equipped with energy filter (slit width 20 eV) and a Gatan K3 camera. The collecting magnification was 26,000 × (3.4 Å/pixel) and the tilt range was −60 ° to 60 ° with 2 ° tilt-series increment and dose-symmetry^55^. The defocus range varied from −7 to −8 μm. The as-obtained tilt series were stacked, aligned and reconstructed into tomograms with *TomoNet*^56^ and *AreTomo*^57^. *MBIR*^58^ was applied to enhance the contrast and alleviate missing wedge problem. Visualization of tomogram slices and three-dimensional rendering of tomograms were done with *IMOD*^59^ and *UCSF ChimeraX*^60^.

### Flagella purification

To purify the extracellular flagella, the *S. oneidensis* cells containing buffer was passed through a 23-gauge syringe needle, following a previously described method^61^. Briefly, cells were resuspended in 6 ml PBS and passed through 26 G 3/8 syringe needle 30 times to shear off the flagella. Cells were then pelleted at 5,000 g for 10 min and resuspended in 6 ml PBS, while the supernatant was retained. The shear-off and centrifugation steps were then repeated twice more, and the combined supernatant was ultracentrifuged (SW41 Ti, 23,000 rpm, 20min, 4 °C) to collect the flagella. The flagella pellet was resuspended in PBS and a final centrifugation (5,000 g, 10 min) was performed to remove unwanted cell debris.

### CryoEM sample preparation and image acquisition

An aliquot of 3 µl of the flagella-containing sample was applied onto Quantifoil holey carbon grids (200 mesh, 2/1) that were glow-discharged for 60 s at 25 mA using the PELCO easiGlow. Using a Vitrobot Mark IV (FEI/Thermo-Fisher) set to 4°C and 100% humidity, excess sample solution on the grids was blotted away by filter paper at a blot force 2 and blot time 4 s. The grids were then plunged into a pre-cooled liquid ethane/propane mixture for vitrification. The plunge-freezing conditions and cell concentration were optimized by screening with an FEI TF20 transmission electron microscope equipped with a Gatan K2 camera.

Single particle cryoEM data were collected as movies on optimized grids in an FEI Titan Krios transmission electron microscope (300 kV) equipped with energy filter (slit width 20 eV) and a Gatan K3 camera at super-resolution mode (81,000 x, 0.55 Å/pixel). The movie was split into 40 frames with a total dose of ∼60 e^-^/Å^2^/micrograph. The defocus range varied from −1.5 to −2.5 μm.

### CryoEM data processing and 3D reconstruction

All frames were performed motion correction and dose weighting using *MotionCor2*^62^. A total of 25,814 drift-corrected micrographs were imported into *Relion 4.0*^63^ and *cryoSPARC*^64^ for further processing.

For the helix reconstruction of the overall filaments, 87.5 k particles manually picked from 3 k representative micrographs, screened by 2D classification, and 58.6 k high-quality particles were selected to train the *Topaz*^65^ for subsequent auto-picking across all micrographs. 912 k particles were obtained from all micrographs via *Topaz* auto-picking. Cylinder density was used as the initial volume and the helix parameters reported in *B. subtilis* were adopted as starting value during helix refinement. Subsequently, 3D classification without alignment was performed to discard junk particles, leaving 464 k good particles. Further CTF refinement accounted for higher-order aberrations, anisotropic magnification, and per-particle defocus, ultimately resolving the overall filament with helical reconstruction to a resolution of around 2.87 Å.

For the asymmetric reconstruction of the curved filaments, we adopted a cryoEM image processing approach similar to that used for *E. coli*^34^. Particles from the helical reconstruction of the overall filament were refined using homogeneous refinement with C1 symmetry to relax the imposed symmetry. The refined particles were then subjected to 3D variability analysis in cryoSPARC, which classified them into 20 clusters. The best cluster, containing approximately 696 k particles and showing clear curvature, was selected for further processing. This included local refinement, followed by CTF refinement and an additional round of local refinement, ultimately yielding a structure of the curved filament at 3.2 Å resolution.

For the asymmetric reconstruction of the hook and HFJ, 14.4k particles (with 80% overlap) and 4.6 k particles were manually picked and imported into cryoSPARC for homogeneous refinement with C1 symmetry, followed by CTF refinement and local refinement. This process ultimately yielded 3D reconstructions of the curved hook and the HFJ at 4.2 Å and 6.1 Å resolution, respectively.

For the reconstruction of the near-junction filament, the particle coordinates were obtained by shifting the refined HFJ particles by 228 pixels in the direction of the filament. Due to the limited number of particles, helical symmetry was applied during reconstruction, followed by CTF refinement and additional helical refinement, which ultimately yielded a structure of the near-junction filament at 3.5 Å resolution.

### Model building and refinement

Model building of all structures started from the predictions of *AlphaFold2*^66^. For FlaA and FlaB in filaments, the whole subunits were initially rigid-body fitted into the helix reconstructed maps of near-junction filament (3.5 Å) and overall filament (2.8 Å) using *UCSF Chimera*^67^. Then the models were refined manually in *COOT*^68^ and further refined with the real space refinement in *Phenix*^36^. For the structures of curved filament maps, the representative FlaB subunit obtained from previous steps was rigid body fitted into the curved map (3.2 Å) and then refined in *Phenix*^36^. FlaA and FlaB features in the four maps above were identified in *COOT*^68^. As for the FlgE in hook (4.2 Å), each domain from *AlphaFold2* prediction was rigid-body fitted separately in *UCSF Chimera*^67^ and then refined in *Phenix*^36^. For the FlgL, FlgK in HFJ (6.1 Å), each domain from *AlphaFold2* prediction was rigid body fitted separately in *UCSF Chimera*^67^.

All visualizations of models and maps in figures were prepared using *UCSF ChimeraX*^60^. Structure alignments were conducted using *PyMOL*^69^ and sequence alignments were presented using *ESPript*^70^.

## Data and code availability

The cryoEM density maps were deposited in the Electron Microscopy Data Bank (EMDB) with accession code: overall flagellar filament (EMD-XXXXX), near-junction filament (EMD-XXXXX), curved filament (EMD-XXXXX), HFJ complex (EMD-XXXXX) and hook (EMD-XXXXX). The corresponding models were deposited in the Protein Data Bank (PDB) with overall flagellar filament (XXXX), near-junction filament (XXXX), and hook (XXXX). This paper does not report the original code. Any additional information required to reanalyze the data reported in this paper is available from the lead contact upon request.

## Acknowledgements

We thank Yao He and Xiaoying Cai for assistance during data acquisition, Yao He and Yun-Tao Liu for advice in data processing. This project is supported by grants from the Noble Family Innovation Fund at CNSI and the National Institutes of Health (R01GM071940 to ZHZ). We acknowledge use of resources in the Electron Imaging Center for Nanomachines supported by UCLA and grants from the NIH (1S10OD018111) and the National Science Foundation (DBI-1338135 and DMR-1548924).

## Author Contributions

Z.H.Z., Y.H. and J.F.M. conceived the project and supervised research; Y.L., Q.L and H.F. prepared samples; Q.L. and H.F. recorded cryoEM and cryoET images and processed the data; Q.L., H.F. and Z.H.Z. interpreted results and wrote the manuscript; all authors reviewed and approved the submitted paper.

