## Supplemental Figures for "Curvature generation and engineering principles from *Shewanella oneidensis* multi-flagellin flagellum"

### Supplementary Figures

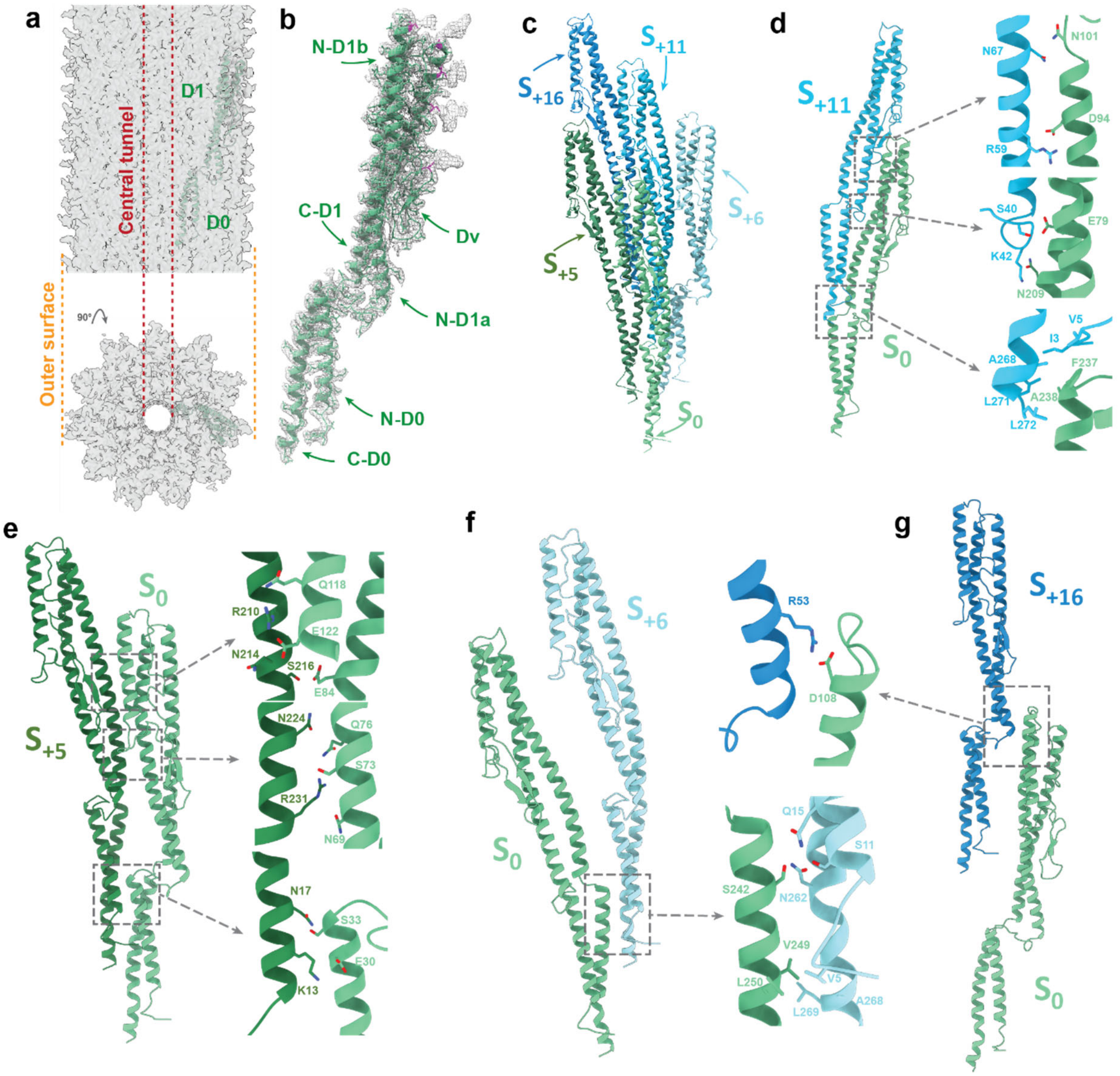


**Figure S1. Structure and arrangement of flagellins. (a)** Side view and top view of the density map (2.8 Å) with a model of FlaB fitted. **(b)** FlaB monomer with corresponding density map. Magenta regions are O-link glycosylation sites. **(c)** Arrangements of subunits in 5-, 6-, 11- and 16-start directions. **(d-g)** Inter-subunit interactions in 11-start direction **(d)**, 5-start direction **(e)**, 6-start direction **(f)** and 16-start direction **(g)**.

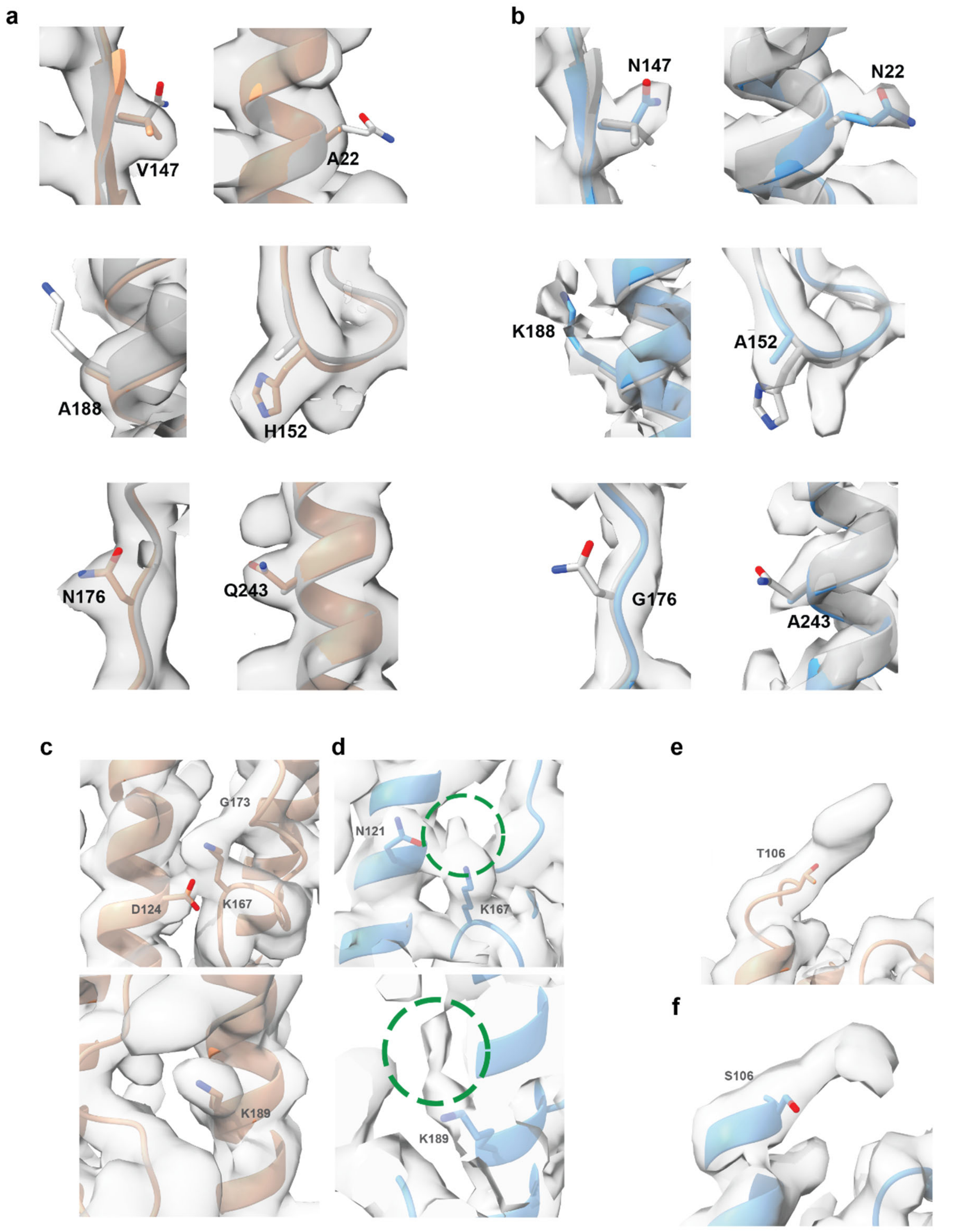


**Figure S2. Structural comparisons between FlaA and FlaB filament. (a)** FlaA (orange) and FlaB (white) models in near-junction filament density map at selected residues. **(b)** FlaA (white) and FlaB (blue) models in overall filament density map at selected residues. **(c-d)** Evidence of the methylation in overall filament density map **(d)** in contrast to near-junction filament density map **(c)**. **(e-f)** O-link glycosylation at residue 106 in near-junction filament density map **(e)** and overall filament density map **(f)**.

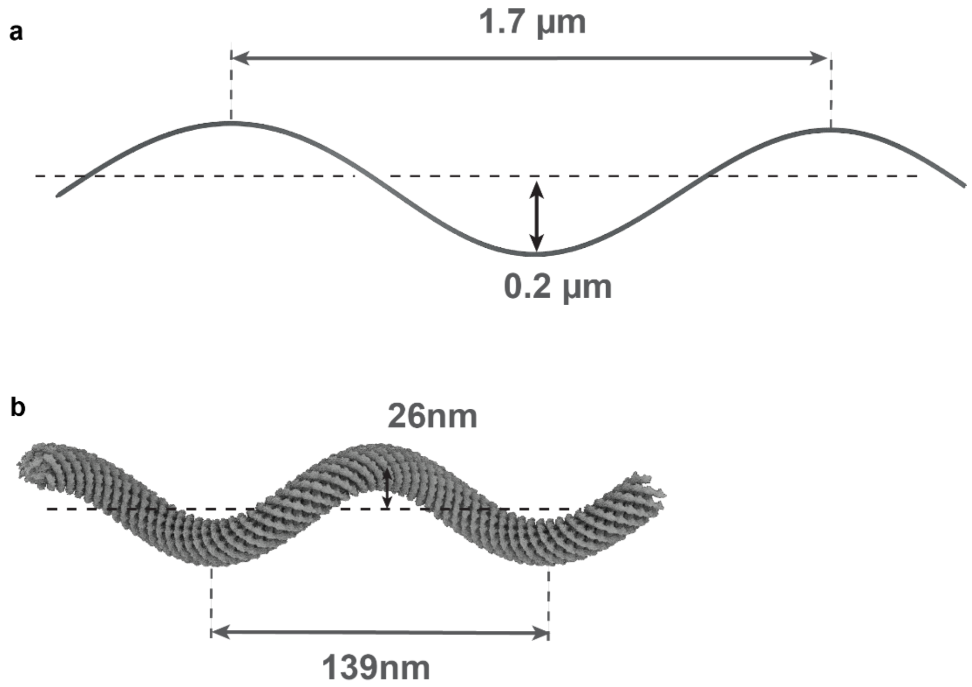


**Figure S3. (a-b)** Supercoils of the flagella filament **(a)** and hook **(b)**.

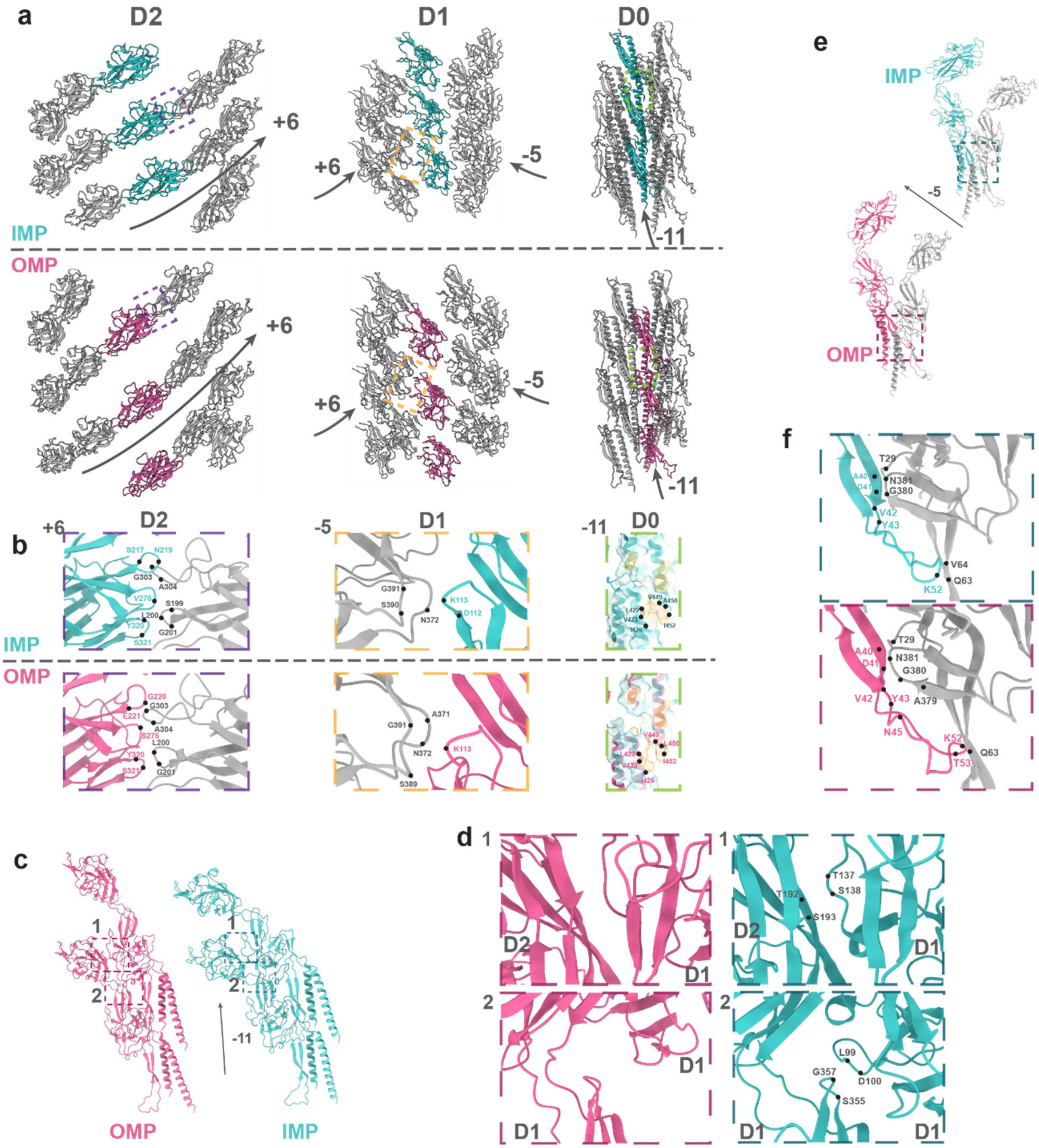


**Figure S4. (a)** Three layers show the arrangements of D2, D1 and D0 domains from IMP and OMP sides. From left to right: D2, D1, D0. Squares indicate the region with interactions. **(b)** Inner-layer interactions in the squared regions of **(a)**. D0 domains are displayed with additional corresponding surface by hydrophobicity. **(c)** Two adjacent subunits in 11-start direction for IMP and OMP. **(d)** D1-D2 and D1-D1 interfaces in the two squared regions of **(c)**. **(e)** Two adjacent subunits in 5-start directions for both IMP and OMP. **(f)** Inter-subunit interactions in the squared regions of **(e)**. The representative residues involved in interactions are marked based on the criteria that Cα distances within 6 Å for potential electrostatic interactions and hydrogen bonds.

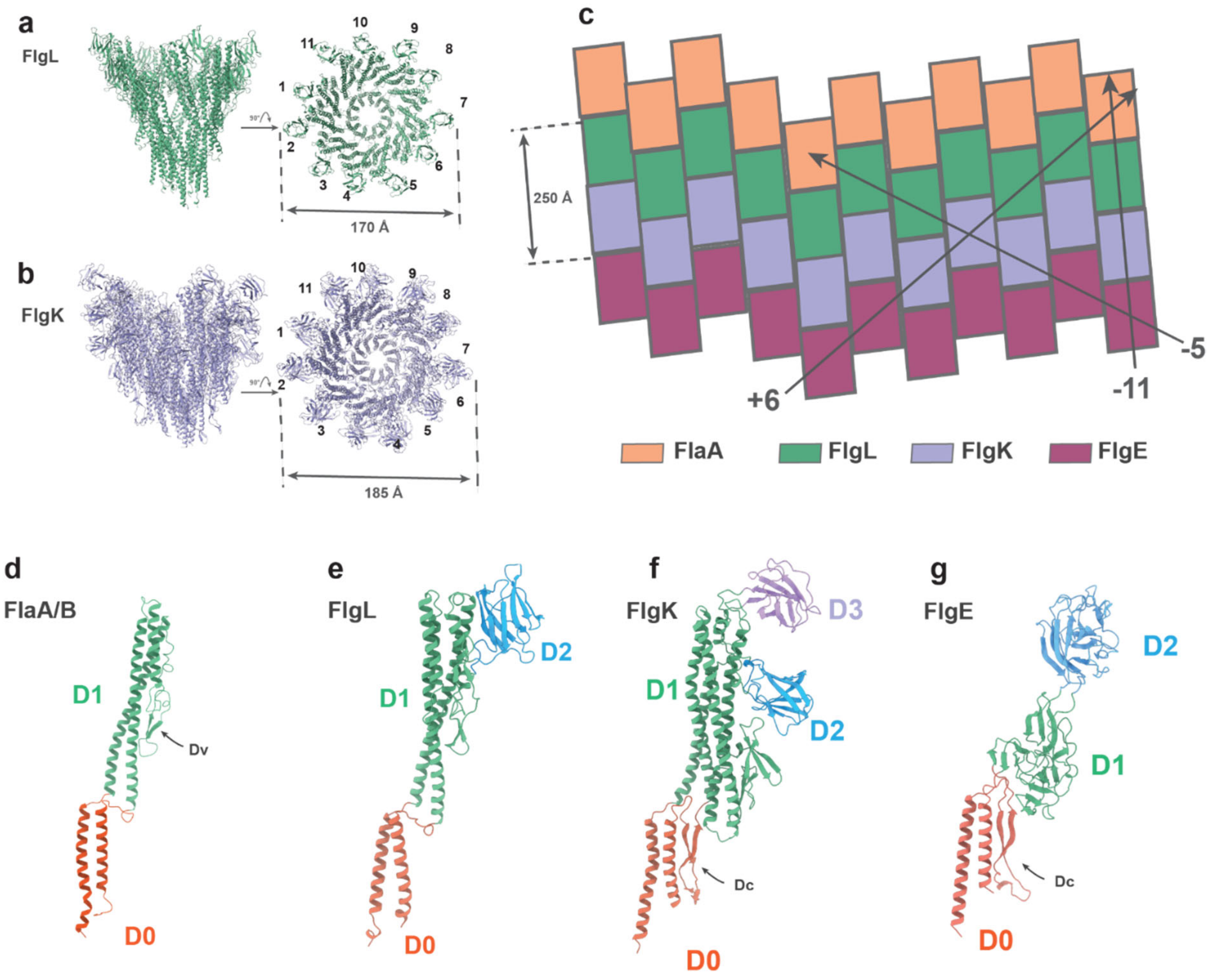


**Figure S5. Arrangement and the quasi-homotypic structures of subunits in junction complex. (a-b)** Side and top views of FlgL **(a)** and FlgK **(b)** helical turns (containing 11 subunits each). **(c)** Schematics of the stacking of different proteins in junction complex. **(d-g)** Model of FlaA/B **(d)**, FlgL **(e)**, FlgK **(f)** and FlgE **(g)** colored differently by domains. Four subunits show gradual transition of structural similarity in common domains D0 and D1. D0 domains are most structurally conserved with two α helices in all subunits, β hairpins occurring in both FlgK (Dc) and FlgE (Dc). In D1 domains, the secondary structure elements of these subunits gradually transit from α helix dominant in FlaA/B to β sheet only in FlgE.

**a**

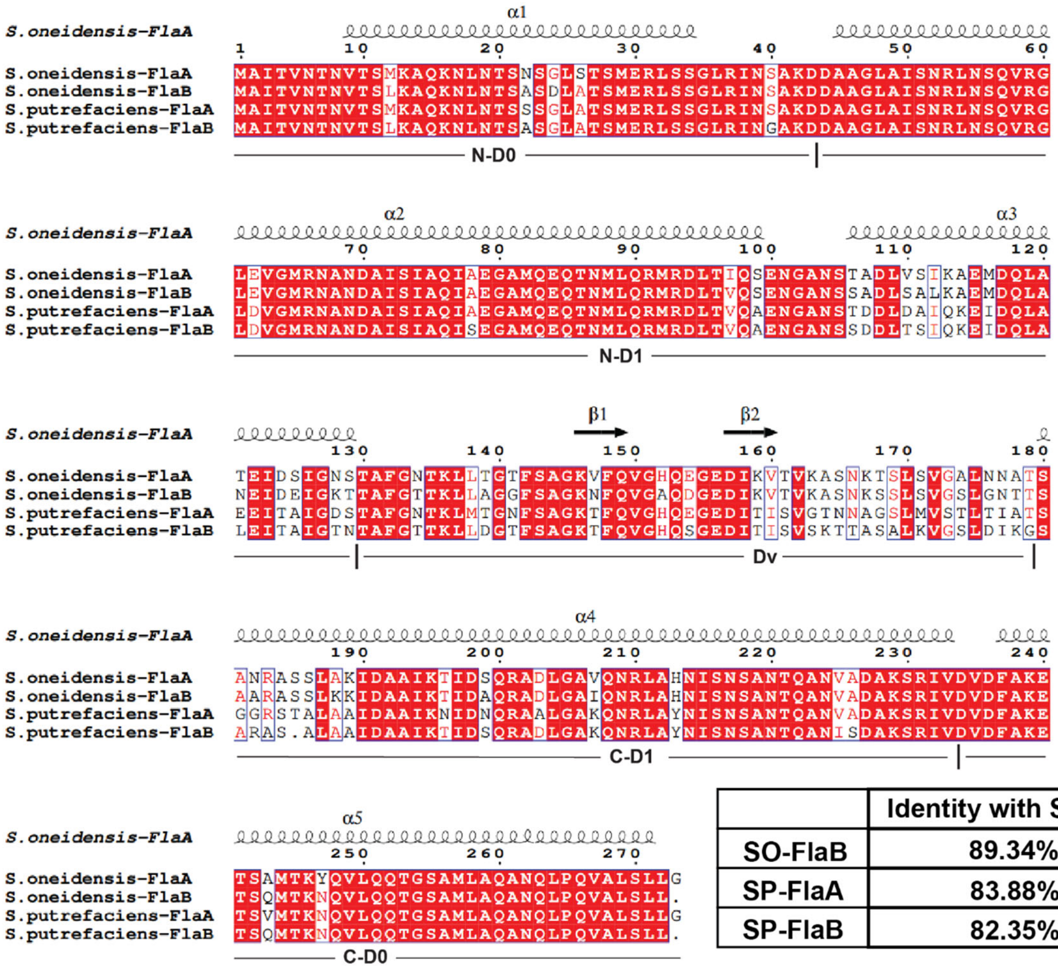


| | Identity with SO-FlaA |
|---|---|
| SO-FlaB | 89.34% |
| SP-FlaA | 83.88% |
| SP-FlaB | 82.35% |

**b**

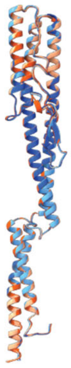


| RMSD/Å | SO-FlaB | SO-FlaA | SP-FlaB |
|---|---|---|---|
| SP-FlaA | 0.46 | 0.45 | 0.53 |
| SP-FlaB | 0.65 | 0.62 | |
| SO-FlaA | 0.31 | | |

**c**

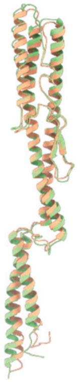


**d**

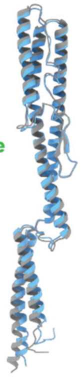


**Figure S6. Conserved structures of flagellins in *S. oneidensis* and *S. putrefaciens*.** **(a)** Sequence alignment of FlaA and FlaB in *S. oneidensis* and *S. putrefaciens*. **(b)** Structural alignment of predicted FlaA and FlaB in *S. oneidensis* and *S. putrefaciens*. **(c)** Structural alignment of predicted FlaA in *S. oneidensis* and its resolved cryoEM structure. **(d)** Structural alignment of predicted FlaB in *S. oneidensis* and its resolved cryoEM structure.
